# Bridging biomolecular modalities for knowledge transfer in bio-language models

**DOI:** 10.1101/2024.10.15.618385

**Authors:** Mangal Prakash, Artem Moskalev, Peter A. DiMaggio, Steven Combs, Tommaso Mansi, Justin Scheer, Rui Liao

## Abstract

In biology, messenger RNA (mRNA) plays a crucial role in gene expression and protein synthesis. Accurate predictive modeling of mRNA properties can greatly enhance our understanding and manipulation of biological processes, leading to advancements in medical and biotechnological applications. Utilizing bio-language foundation models allows for leveraging large-scale pretrained knowledge, which can significantly improve the efficiency and accuracy of these predictions. However, mRNA specific foundation models are notably limited posing challenges for efficient predictive modeling in mRNA-focused tasks. In contrast, DNA and protein modalities have numerous general-purpose foundation models trained on billions of sequences. This paper explores the potential for adaptation of existing DNA and protein bio-language models for mRNA-focused tasks. Through experiments using various mRNA datasets curated from both public domain and internal proprietary database, we demonstrate that pre-trained DNA and protein models can be effectively transferred for mRNA-focused tasks using various adaptation techniques such as probing, full-rank, and low-rank finetuning. In addition, we identify key factors that influence successful adaptation, offering guidelines on when general-purpose DNA and protein models are likely to perform well for mRNA-focused tasks. We further assess the impact of model size on adaptation efficacy, finding that medium-scale models often outperform larger ones for cross-modal knowledge transfer. We conclude that by leveraging the interconnectedness of DNA, mRNA, and proteins, as outlined by the central dogma of molecular biology, the knowledge in foundation models can be effectively transferred across modalities, significantly enhancing the repertoire of computational tools available for mRNA analysis.

## 1 Introduction

Messenger-RNA (mRNA) is central to molecular biology, serving as the template for protein synthesis and thereby influencing virtually every cellular process. Developing computational models that can robustly analyze mRNA data is essential for advancing our understanding of gene regulation and enhancing our ability to engineer biological systems.

The powerful method of analyzing molecular biology sequences such as RNA, DNA, or protein involves the use of *Language Models (LMs)* to extract informative representations (Zhou et al., 2023; Dalla-Torre et al., 2023; Nguyen et al., 2024b,a; Elnaggar et al., 2021, 2023; Lin et al., 2023; Nijkamp et al., 2023; Ruffolo et al., 2021). These models have demonstrated substantial success across various biological applications, ranging from prediction of protein structures and properties, understanding of genetic variations, to decoding of complex genetic information.

Unlike DNA and protein modalities, which have numerous general-purpose bio-LMs trained on hundreds of millions to billions of sequences, publicly available mRNA models remain underrepresented. Although a few RNA language models are publicly available, they are pretrained on much smaller datasets (approximately 10-fold smaller) and are tailored to specific RNA subtypes or regions. This limited scale, combined with the lack of specificity to mRNA regions, restricts the generalizability of these models for comprehensive mRNA analysis. The development of robust mRNA models is further hindered by the scarcity of high-quality, curated mRNA datasets and the high dimensionality of mRNA sequences, which can reach up to 100k nucleotides.

Recognizing these challenges, we aim to explore the adaptation of existing DNA and protein LMs for downstream mRNA-focused tasks. Although DNA, proteins, and mRNA each possess unique biological roles and properties, they are interconnected by the central dogma of molecular biology (Figure 1). This connection provides a unified flow of genetic information across these biomolecular modalities and serves as a solid basis for knowledge transfer. In this paper, we aim to unveil the knowledge transfer enabled by the central dogma to harness the extensive pretraining of DNA and protein LMs for mRNA-focused tasks. To this end, we explore several adaptation techniques, including probing, full-rank, and low-rank finetuning. Our findings reveal that general DNA and protein models not only outperform the traditional one-hot mRNA-level baselines but often also surpass general-purpose RNA-specific LMs when applied to mRNA-focused tasks. Moreover, we uncover several key factors impacting successful adaptation and provide guidelines on when general-purpose DNA and protein models can be expected to demonstrate good adaptation performance.

**Figure 1:**
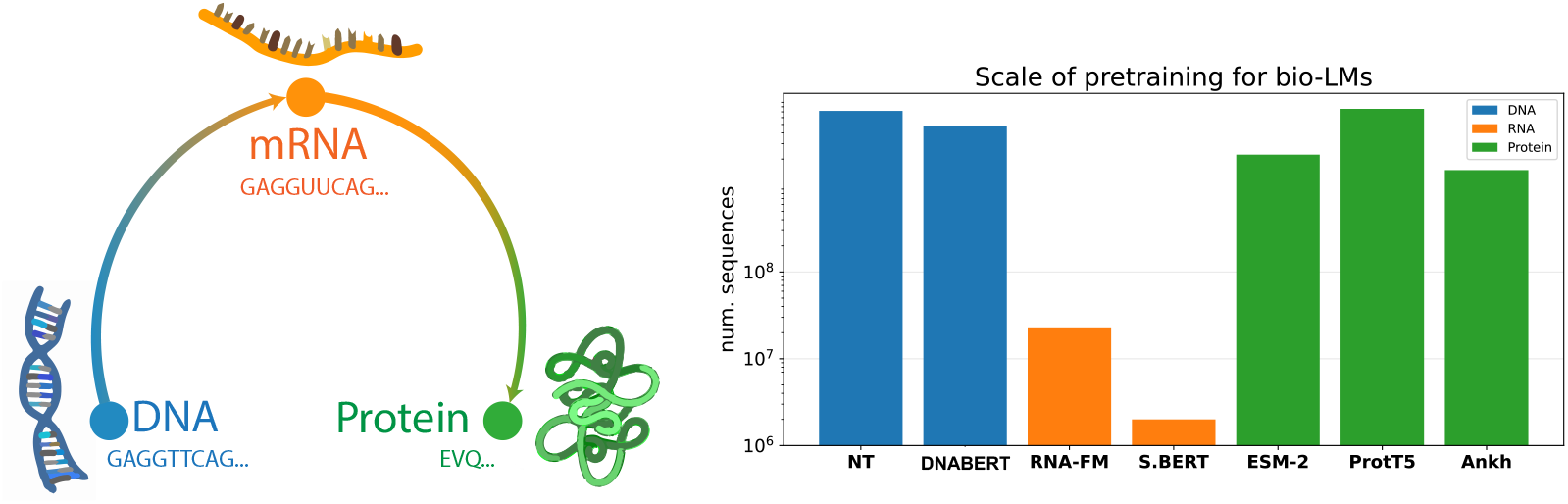
**Left:** Central dogma of molecular biology. DNA is transcribed into messanger-RNA (mRNA) which is then translated to protein. **Right:** Comparing scale of data used for pre-training DNA, RNA, and protein language models. Public RNA language models are trained on datasets with several magnitudes smaller than ones used for pretraining DNA or protein models.

To sum up, we make the following contributions:

- **Cross-modal adaptation of bio-language models:** We show that DNA and protein LMs can be effectively adapted for mRNA-focused tasks under various adaptation strategies, including full finetuning, low-rank finetuning, and probing. This significantly enriches the computational toolkit available for mRNA analysis.
- **Factors influencing cross-modal adaptation efficiency:** We delve into several datasets, from both public domain and internal proprietary database, and analyze the influencing factors that affect the efficiency of adapting protein and DNA LMs for mRNA-focused tasks, enhancing understanding of cross-modal adaptation dynamics.
- **Impact of model size:** Through comprehensive testing on various publicly available and internally acquired mRNA datasets, we identify model size as a key factor that influences the performance of DNA and protein LMs in cross-modal knowledge transfer.

## 2 Related work

### Language models for DNA, RNA, and Protein

Transformer-based LMs, structured state space LMs and long-convolutional LMs have become state-of-the-art in modeling bio-languages. The standard approach involves pretraining a LM on a large database with masked language or causal masking objectives. This enables learning the general grammar of the bio-language of interest, facilitating fast adaptation for various downstream tasks. For DNA modality, transformer-based models such as DNABERT (Ji et al., 2021), DNABERT-2 (Zhou et al., 2023), Nucleotide Transformer (Dalla-Torre et al., 2023), and long-convolutional models like HyenaDNA (Nguyen et al., 2024b) and Evo (Nguyen et al., 2024a) have demonstrated excellent performance in genomics applications. For protein modality, LMs such as those from ESM-family (Lin et al., 2023), ProtTrans (Elnaggar et al., 2021), ProGen2 (Nijkamp et al., 2023), Ankh (Elnaggar et al., 2023), and ProtMamba (Sgarbossa et al., 2024) have excelled in predicting protein structure and functions for various protein families. The data scale for pretraining such DNA and protein LMs typically varies from 100M to 2B sequences. Additionally, most of these bio-LMs are openly accessible through platforms like HuggingFace (Wolf et al., 2019), facilitating their rapid off-the-shelf deployment.

In contrast to this, publicly available general-purpose RNA modality LMs are pretrained on much smaller datasets and often are limited to specific RNA regions and species due to the scarcity of suitable RNA datasets. For instance, RNA-FM (Chen et al., 2022) is pretrained on 23 million non-coding RNAs, UTR-LM (Chu et al., 2024) and UTRBERT (Yang et al., 2023) are pretrained with 700K and 20K 5’ and 3’ UTR-RNA sequences respectively while SpliceBERT (Chen et al., 2023) is pretrained on 2 million precursor mRNA sequences. Proprietary RNA modality LMs such as those proposed by Li et al. (2023), Celaj et al. (2023), and Wang et al. (2023) are not accessible under permissible licenses. There are currently no bio-LMs specifically designed for mRNA that are freely accessible and permissibly usable.

### Knowledge transfer in language models

Knowledge transfer in LMs has become critical for leveraging information from data-rich domains to improve performance in resource-constrained areas. This approach has gained prominence with large pretrained models like BERT (Devlin et al., 2018) and GPT (Brown et al., 2020). Techniques such as fine-tuning (Howard and Ruder, 2018), distillation (Hinton et al., 2015), and parameter-efficient methods like adapters (Houlsby et al., 2019; Hu et al., 2021) and prompt-tuning (Lester et al., 2021) have shown success in transferring knowledge across diverse tasks and domains. These methods often outperform models trained from scratch on limited domain-specific data. Our work extends this paradigm to biological domains, exploring knowledge transfer between different molecular modalities to address the challenges of limited mRNA-specific data and models.

## 3 Method

### 3.1 Models and datasets

#### Choosing bio-language models

We employ state-of-the-art transformer-based bio-LMs specializing in different biomolecular modalities and recognized for their superlative performance. As a DNA model, we employ the Nucleotide Transformer (NT) (Dalla-Torre et al., 2023). For proteins, we use ESM-2 (Lin et al., 2023). Notably, both NT and ESM-2 offer various model sizes, allowing us to study the impact of model scaling on cross-modal knowledge transfer.

Given that DNA and protein LMs use different vocabularies compared to mRNA, adapting these models for mRNA-focused tasks first requires addressing this difference. Our approach leverages the well-established central dogma of molecular biology, which outlines the flow of genetic information from DNA to mRNA to protein. Specifically, mRNA sequences with known start and stop codons can be mapped back to their corresponding DNA and protein sequences. This conversion process involves two key steps: converting mRNA to DNA through reverse transcription, where nucleotide uracil (*U*) is replaced by nucleotide thymine (*T*) (Temin and Mizutami, 1970; Baltimore, 1970), and translating mRNA codons (sequences of three nucleotides) into amino acids to form proteins through translation (Chaffey, 2003). Thus, we adapt mRNA sequences for inputs to NT and ESM-2 mirroring the natural biological processes of reverse transcription and translation. Although these biological principles are well-established, they have not been previously utilized in the context of adapting DNA and protein LMs for mRNA-focused tasks.

Since no mRNA specific model is publicly available under permissible license, we employ RNA-FM (Chen et al., 2022) and SpliceBERT (Chen et al., 2023) as two in-modality baselines. RNA-FM, despite being pretrained on non-coding RNAs (ncRNAs), has been acknowledged as the strongest baseline in recent studies (Franke et al., 2024; Nguyen et al., 2024a) for various general purpose RNA-focused tasks and used in comparison with proprietary models such as those by Boyd et al. (2023); Li et al. (2023). In comparison, SpliceBERT has been pretrained on smaller pre-mRNA dataset but is the closest available LM to mRNA modality from evolutionary perspective.

Unlike RNA-FM, the other models are accessible via HuggingFace, facilitating easy modification and finetuning.

#### Datasets

Since existing RNA models have not been pretrained and evaluated for mRNA specifically, their associated downstream datasets and tasks cannot be used for our study that is specifically designed for mRNA-focused tasks. Instead, we curate 4 public mRNA datasets and also acquire 2 internal proprietary mRNA datasets. The considered datasets are diverse spanning multiple species and covering various mRNA-focused downstream tasks:

- *E. coli* dataset (Ding et al., 2022): contains 4450 mRNA sequences along with experimental data binning of mRNA to protein expression within 6 classes.
- iCodon stability dataset (Diez et al., 2022): includes 1144 mRNA sequences with thermostability profiles from humans, mice, frogs, and fish.
- SARS-CoV-2 Vaccine Degradation dataset (Leppek et al., 2022): comprises a collection of 4893 mRNA sequences with degradation rates as labels per nucleotide for the first 68 nucleotides.
- Fungal expression dataset (Wint et al., 2022): consists of 3140 mRNA sequences with mRNA to protein yield labels.
- Internal propreitary Antibody mRNA (Ab1) expression dataset: contains 1200 mRNA sequences with mRNA to protein expression labels.
- Internal propreitary Antibody mRNA (Ab2) expression dataset: contains 3442 mRNA sequences with mRNA to protein expression labels.

We preprocess all mRNA sequences, removing invalid ones based on the following criteria: (i) sequence is divisible by 3 in length^2^, (ii) sequence begins with start codon *AUG* and ends with one of the stop codons *UAA, UAG*, or *UGA*, (iii) sequence contains only *A, U, G, C, N* nucleotides. These criteria are suitable for all datasets except the SARS-CoV-2 dataset, which lacks clear start and stop codons, creating ambiguity in translation to protein for ESM-2. This affects the performance of ESM-2 as discussed in Section 4.4. Additionally, sequence lengths are limited to 1024 tokens owing to the limitation imposed by both the RNA models (RNA-FM and SpliceBERT).

All tasks except for *E. coli* expression prediction are regression tasks. We used a 70 : 15 : 15 random split for training, validation, and testing.

### 3.2 Adaptation strategies for knowledge transfer

#### Probing

We first evaluate knowledge transfer by probing the quality of DNA and protein LM embeddings for mRNA tasks. We freeze the LM, extract embeddings from the last layer, and train a downstream head to map these embeddings to task-specific labels (see Figure 2 top left panel). We compare TextCNN (Chen, 2015) and LSTM (Hochreiter and Schmidhuber, 1997) heads, finding TextCNN to perform better or on par with LSTM for most datasets (details in Appendix A). Consequently, we use TextCNN head for all experiments. This approach allows us to assess the transferability of learned representations without modifying the original backbone LMs.

**Figure 2:**
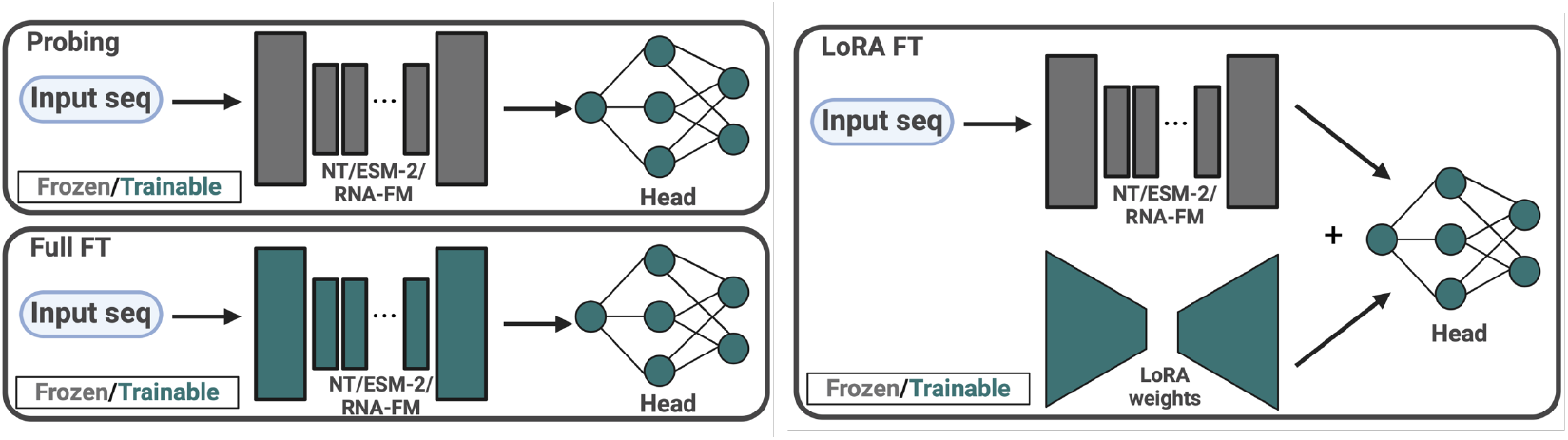
Overview of the adaptation strategies used to evaluate knowledge transfer and fine-tuning in bio-LMs for mRNA tasks. The top left panel illustrates the probing approach, where embeddings are extracted from the frozen DNA or protein LM and mapped to task-specific labels using a TextCNN head. The bottom left panel shows full supervised fine-tuning (Full FT), where all parameters of the model backbone and head are updated. The right panel depicts parameter-efficient fine-tuning using Low-Rank Adaptation (LoRA FT), where only specific layers or parameters are adapted while the rest of the model remains unchanged.

#### Supervised finetuning

We also explore supervised finetuning of the bio-LMs on each labeled dataset. We evaluate two approaches: full finetuning (Full FT), where all parameters of the model backbone and head are updated (see Figure 2 bottom left panel), and parameter-efficient finetuning using Low-Rank Adaptation (Hu et al., 2021) (LoRA FT) (see Figure 2 right panel). This comparison helps us understand the trade-offs between comprehensive model updating and more efficient, targeted adaptation strategies.

### 3.3 Implementation details

#### Training and hyperparameters

For probing experiments, we use AdamW optimizer (Kingma and Ba, 2014). The batch size and head learning rate are determined through a grid search on the validation set, exploring batch sizes of 16, 32, 64, and 128, and learning rates of 1 *×* 10^*−*4^, 3 *×* 10^*−*4^, 5 *×* 10^*−*4^, 7 *×* 10^*−*4^, and 9 *×* 10^*−*4^. For full finetuning, we perform a grid search over batch sizes and head learning rates in the same range as for probing experiments to select the hyperparameters. Additionally, we consistently use a backbone learning rate of 5 *×* 10^*−*5^ during finetuning, as changing this rate always leads to performance deterioration in all experiments. For LoRA FT, we additionally perform a grid search over LoRA ranks 16, 32, 64, and 128 and alpha values 0.25*×*, 0.5*×*, 1*×*, 2*×* the chosen ranks for each dataset.

For probing and finetuning, we select the 100*M* parameter NT model and the 150*M* million parameter ESM-2 model as these are closest in size to the available 100*M* parameter RNA-FM model. SpliceBERT, on the other hand, is only available at 20*M* parameter scale but is pretrained on pre-mRNA data which is evolutionarily closer to mRNA modality we are focused on.

## 4 Experiments

### 4.1 Cross-modal adaptation of DNA and protein bio-language models to mRNA-focused tasks

Table 1 displays the performance of different bio-LMs using various adaptation techniques. We compare against traditional one-hot baselines on DNA, RNA, and protein levels, where only the head is trained with one-hot embeddings as is typical in prior works (Harmalkar et al., 2023; Boyd et al., 2023).

**Table 1:**
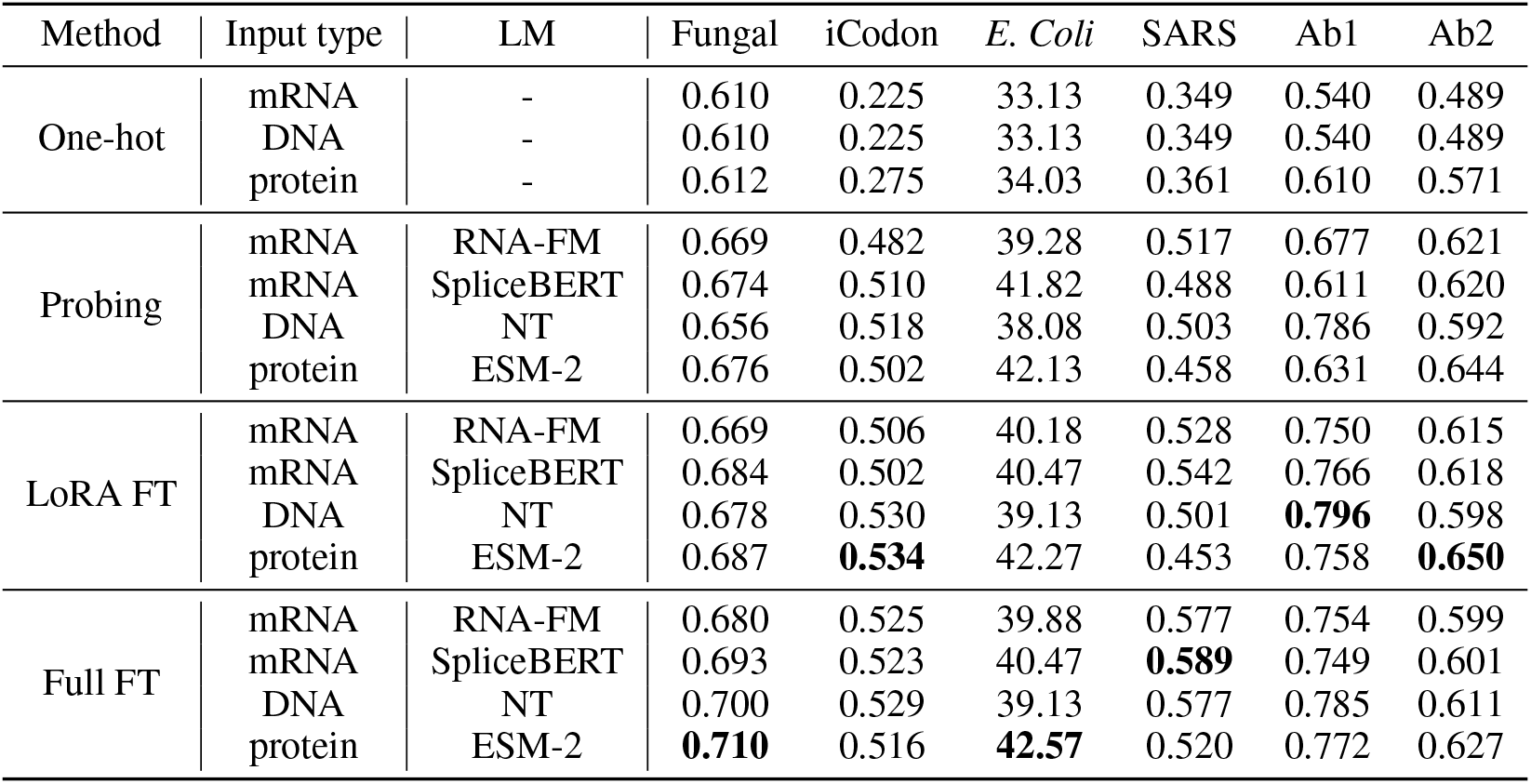
Both DNA and protein LMs adapted for mRNA-focused tasks perform well. Spearman correlation is reported for Fungal, iCodon, SARS, Ab1, and Ab2 datasets, while classification accuracy is reported for *E. Coli* dataset. Bold indicates the best performing method. Protein or DNA LMs perform best in 5 out of 6 datasets with either full FT or parameter-efficient LoRA FT. This demonstrates that DNA and protein language models can serve as powerful alternatives to RNA-specific models for mRNA analysis, owing to their extensive pretraining on larger datasets and the interconnectedness of biomolecular modalities within the central dogma paradigm.

| Method | Input type | LM | Fungal | iCodon | *E. Coli* | SARS | Ab1 | Ab2 |
| --- | --- | --- | --- | --- | --- | --- | --- | --- |
| One-hot | mRNA | - | 0.610 | 0.225 | 33.13 | 0.349 | 0.540 | 0.489 |
|  | DNA | - | 0.610 | 0.225 | 33.13 | 0.349 | 0.540 | 0.489 |
|  | protein | - | 0.612 | 0.275 | 34.03 | 0.361 | 0.610 | 0.571 |
| Probing | mRNA | RNA-FM | 0.669 | 0.482 | 39.28 | 0.517 | 0.677 | 0.621 |
|  | mRNA | SpliceBERT | 0.674 | 0.510 | 41.82 | 0.488 | 0.611 | 0.620 |
|  | DNA | NT | 0.656 | 0.518 | 38.08 | 0.503 | 0.786 | 0.592 |
|  | protein | ESM-2 | 0.676 | 0.502 | 42.13 | 0.458 | 0.631 | 0.644 |
| LoRA FT | mRNA | RNA-FM | 0.669 | 0.506 | 40.18 | 0.528 | 0.750 | 0.615 |
|  | mRNA | SpliceBERT | 0.684 | 0.502 | 40.47 | 0.542 | 0.766 | 0.618 |
|  | DNA | NT | 0.678 | 0.530 | 39.13 | 0.501 | **0.796** | 0.598 |
|  | protein | ESM-2 | 0.687 | **0.534** | 42.27 | 0.453 | 0.758 | **0.650** |
| Full FT | mRNA | RNA-FM | 0.680 | 0.525 | 39.88 | 0.577 | 0.754 | 0.599 |
|  | mRNA | SpliceBERT | 0.693 | 0.523 | 40.47 | **0.589** | 0.749 | 0.601 |
|  | DNA | NT | 0.700 | 0.529 | 39.13 | 0.577 | 0.785 | 0.611 |
|  | protein | ESM-2 | **0.710** | 0.516 | **42.57** | 0.520 | 0.772 | 0.627 |

All four evaluated bio-LMs significantly outperform their respective modality specific one-hot baselines across all datasets. Despite being pretrained on different modalities, NT and ESM-2 show promising performance for mRNA-focused tasks. In probing experiments, ESM-2 and NT together outperform the RNA-specific RNA-FM on 5 out of 6 datasets while outperforming SpliceBERT on 3 of the datasets and showing on-par performance on the rest.

Full and low-rank finetuning further improve performance over probing-based approaches. With either full or LoRA finetuning, protein and DNA LMs surpass both RNA-FM and SpliceBERT on 5 out of 6 datasets. ESM-2 achieves the best performance on 4 datasets while NT has the best performance on 1 dataset.

*This suggests that DNA and protein language models can serve as powerful alternatives to RNA-specific models for mRNA analysis*. We can attribute this success of protein and DNA LMs to two factors: (1) the extensive pre-training of these models on larger datasets compared to RNA models and (2) the interconnectedness of biomolecular modalities within the central dogma paradigm, which facilitates knowledge transfer between them. In addition, we identify several factors that impact successful adaptation and discuss them in detail in the following subsections.

### 4.2 Influence of model size on cross-modal knowledge transfer

Since NT and ESM-2 are available in different sizes, we also explore the effect of model sizes on cross-modality knowledge transfer. While scaling laws in NLP and biomolecular domains often suggest that larger models capture more complex dependencies (Kaplan et al., 2020; Dalla-Torre et al., 2023; Lin et al., 2023), recent findings also highlight *inverse scaling* (McKenzie et al., 2023), where larger models do not uniformly outperform smaller ones on all tasks.

Our probing results in Table 2 and Figure 3 for various sizes of ESM-2 and NT models reveal that increasing model size does not monotonically improve performance in cross-modal adaptation for bio-languages. For both NT and ESM-2 models, performance initially improves with increasing model sizes but deteriorates beyond a certain point, except for the Fungal dataset.

**Table 2:**
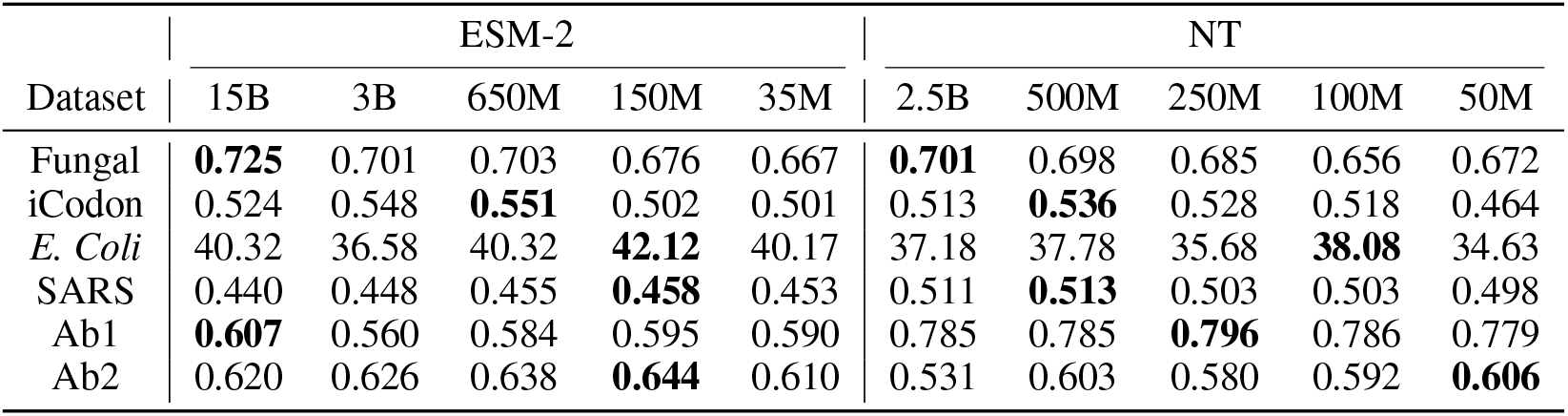
Probing performance across various sizes for ESM-2 and NT models. Bold indicates best performing model. Larger ESM-2 and NT models show diminished performance beyond a certain size, indicating challenges in adapting to mRNA tasks. This may result from retaining pretrained knowledge, especially with smaller downstream datasets. These results highlight the importance of carefully selecting model size for mRNA-focused bio-LM adaptation.

| Dataset | ESM-2 |  |  |  |  | NT |  |  |  |  |
| --- | --- | --- | --- | --- | --- | --- | --- | --- | --- | --- |
|  | 15B | 3B | 650M | 150M | 35M | 2.5B | 500M | 250M | 100M | 50M |
| Fungal | **0.725** | 0.701 | 0.703 | 0.676 | 0.667 | **0.701** | 0.698 | 0.685 | 0.656 | 0.672 |
| iCodon | 0.524 | 0.548 | **0.551** | 0.502 | 0.501 | 0.513 | **0.536** | 0.528 | 0.518 | 0.464 |
| *E. Coli* | 40.32 | 36.58 | 40.32 | **42.12** | 40.17 | 37.18 | 37.78 | 35.68 | **38.08** | 34.63 |
| SARS | 0.440 | 0.448 | 0.455 | **0.458** | 0.453 | 0.511 | **0.513** | 0.503 | 0.503 | 0.498 |
| Ab1 | **0.607** | 0.560 | 0.584 | 0.595 | 0.590 | 0.785 | 0.785 | **0.796** | 0.786 | 0.779 |
| Ab2 | 0.620 | 0.626 | 0.638 | **0.644** | 0.610 | 0.531 | 0.603 | 0.580 | 0.592 | **0.606** |

**Figure 3:**
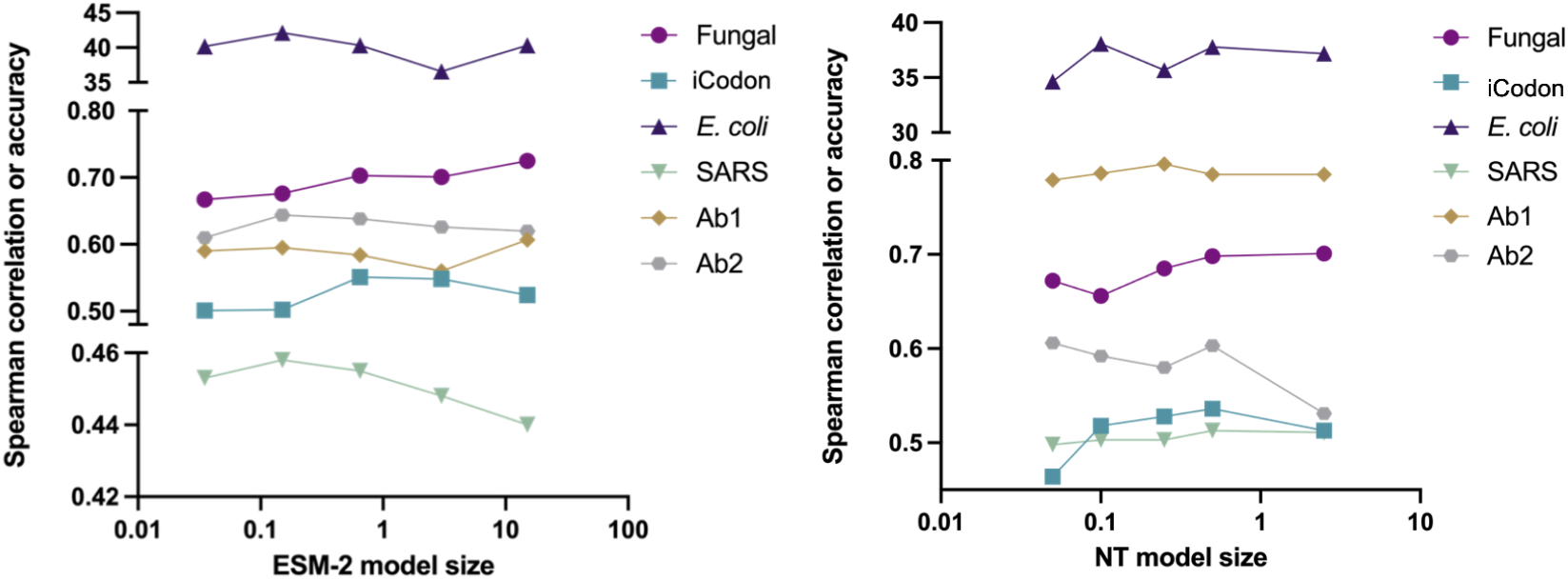
Performance analysis of different sizes (in billions of parameters) of ESM-2 (left) and NT (right) in cross-modal adaptation for mRNA-focused tasks. Larger models tend to exhibit poorer performance beyond a certain size for both ESM-2 and NT. This suggests that larger DNA and protein models may face challenges in adapting to mRNA tasks, potentially due to the preservation of their pretrained knowledge, particularly when the downstream dataset is limited in scale. These findings underscore the need to carefully consider model size when adapting bio-LMs for mRNA-focused applications.

This behavior suggests that larger DNA and protein models may struggle to adapt to mRNA-focused tasks in the cross-modal knowledge transfer setting. This may be attributed to the tendency of large models to preserve their inertia, especially when the scale of the downstream dataset is limited for finetuning (McKenzie et al., 2023). Our results highlight the importance of carefully considering model size when adapting DNA and protein LMs for mRNA-focused applications.

### 4.3 Impact of nature of downstream task

ESM-2 performs exceptionally well in iCodon stability and expression prediction tasks for datasets such as fungal, E. Coli, Ab1, and Ab2. This success can be attributed to ESM-2’s pretraining on extensive protein sequence databases. It is well established that protein sequences encode thermal stability-related information (Teng et al., 2010). Moreover, thermal stability is known to correlate with protein expression levels, as more stable proteins are typically expressed at higher levels (Hanson et al., 2019; Bæk et al., 2023). Given this biological context, ESM-2’s pretraining on diverse and extensive protein datasets enables it to capture and leverage thermal stability information inherent in protein sequences. The model’s ability to learn these nuanced biological relationships from its pretraining data translates into superior performance on tasks that require understanding of protein stability and its impact on expression.

### 4.4 Effect of dataset quality on cross-modal knowledge transfer

Although degradation also correlates with thermal stability (Simantov and Goyal, 2022), still ESM-2 exhibits the poorest performance among all models on the SARS dataset (Leppek et al., 2022) where the objective is to predict degradation per nucleotide. This discrepancy can be attributed to the inherent nature of protein models, which operate at the amino acid level and consequently encounter a loss of resolution when predicting nucleotide level properties. Furthermore, unlike other datasets, SARS dataset used for this task lacks a well-defined start and stop codon for some sequences, further compromising the accuracy of the translation of mRNA sequences performed for modeling purposes. Consequently, the resulting protein sequences used for modeling are noisy. Therefore, in cases where the focus is on predicting nucleotide-level degradation, ESM-2 proves to be less accurate due to its inherent amino acid-centric context and the limitations imposed by dataset quality. On the other hand, for this task both RNA-FM and NT perform equally well owing to their ability to reason at the nucleotide level.

## 5 Discussion and conclusion

Our study demonstrates the successful adaptation of DNA and protein LMs for mRNA tasks. While direct comparison of these models is complex due to diverse pretraining contexts, our focus was on exploring whether easily accessible protein and DNA models can facilitate downstream tasks in underexplored modalities like mRNA, where specialized LMs are non-existent.

The results reveal significant cross-modal adaptation potential, with protein and DNA LMs showing promise for mRNA analysis and offering viable alternatives. These findings can also serve as a benchmark for assessing newly pretrained mRNA LMs.

Interestingly, we also observed that larger models do not always perform better in cross-modal adaptation, suggesting the need for careful model selection based on the specific task and available mRNA data. Furthermore, we identified other important factors affecting adaptation success of DNA and protein LMs: the nature of downstream tasks and the quality of available data.

These findings highlight the immense potential as well as pitfalls of leveraging knowledge transfer from existing DNA and protein LMs for mRNA related tasks. Future work could explore more sophisticated adaptation techniques and investigate the integration of multiple modalities to further enhance mRNA analysis capabilities.

## A Appendix

### Downstream head networks

We experiment with two head architectures: TextCNN (Chen, 2015) and LSTM (Hochreiter and Schmidhuber, 1997), each with approximately 1.2 million parameters. The TextCNN model projects input features into a 640-dimensional space using a linear layer, and passes them through 3 convolutional layers with kernel sizes of 3, 4, 5 and channel dimension of 100, followed by ReLU activation. Max-pooling is applied across the temporal dimension, resulting in a feature map of size 100 for each kernel size. These are concatenated, subjected to 0.2 dropout, and fed into a fully connected layer for final predictions. The LSTM head processes input sequences through an LSTM layer with a hidden dimension of 640. The output undergoes max-pooling across the sequence length, followed by fully connected layers. Dropout of 0.2 is applied before the final fully connected layer.

#### A.1 Head network ablation

**Table 3:** Comparison of TextCNN and LSTM heads for probing experiments. TextCNN head performs best with almost all models across all datasets.

|  |  | Fungal | iCodon | *E. Coli* | SARS | Ab1 | Ab2 |
| --- | --- | --- | --- | --- | --- | --- | --- |
| LSTM | RNA-FM | 0.646 | 0.420 | 40.02 | 0.509 | 0.630 | 0.628 |
|  | NT | 0.615 | 0.485 | 32.68 | 0.493 | 0.772 | 0.593 |
|  | ESM-2 | 0.682 | 0.460 | 38.68 | 0.456 | 0.688 | 0.638 |
| TextCNN | RNA-FM | 0.669 | 0.482 | 39.280 | 0.517 | 0.677 | 0.621 |
|  | NT | 0.656 | 0.518 | 38.081 | 0.503 | 0.786 | 0.592 |
|  | ESM-2 | 0.676 | 0.502 | 42.128 | 0.458 | 0.631 | 0.644 |

## Footnotes

Foundation Models for Science Workshop,38th Conference on Neural Information Processing Systems (NeurIPS 2024).

2 to ensure proper mRNA to protein translation for ESM-2

